# Consolation behaviour in pigs: Prior exposure to group members in need of help drives targeted affiliation and facilitates social buffering

**DOI:** 10.64898/2026.04.02.716034

**Authors:** Alvaro Lopez Caicoya, Wiktoria Janicka, Liza R. Moscovice

**Affiliations:** Psychophysiology Working Group, Research Institute for Farm Animal Biology (FBN), 18196 Dummerstorf, Mecklenburg-Vorpommern, Germany; Institute of Biological Basis of Animal Production, Faculty of Animal Sciences and Bioeconomy, University of Life Sciences in Lublin, Lublin, Poland

## Abstract

We assessed whether pigs provide consolation, referring to targeted affiliation that attenuates a partner’s stress, under experimental conditions that manipulated exposure to stressed partners. Using a within-subject design, 74 pigs were tested in three contexts: a helping task in which group members could observe and help a trapped focal pig to return to the group, a direct-reunion, in which group members were naïve to the experience of a separated focal pig until reunion, and an undisturbed control. We measured affiliative and non-affiliative interactions, anxiety behaviours and changes in salivary cortisol. Only the helping context satisfied most consolation criteria: there were selective increases in unidirectional affiliative contacts from the observer to the focal pig, non-affiliative interactions remained at baseline, and focal pigs showed fewer anxiety behaviours. In contrast, direct-reunions triggered increases in affiliative and non-affiliative interactions and higher anxiety. Cortisol increased during both direct-reunions and helping, but its level was not linked to affiliation. Results add to growing evidence for consolation behaviour in pigs and suggest best practices for reintegrating pigs into groups. Graded reintroductions that allow observers to assess the emotional state of targets may promote social buffering, whereas abrupt regrouping may trigger more generalized arousal or personal distress.

## 1. Introduction

### 1.1 Comparative evidence for consolation

In humans, greater levels of social support, defined as the experience or perception that one is cared for by others (Taylor, 2011), are related to a range of positive physical and mental health outcomes (Vila, 2021). One of the main mechanisms by which social support is believed to have positive health effects is through the provision of advice, aid and/or emotional support that helps individuals to better cope with stressful events (Taylor, 2011) and to avoid detrimental impacts of prolonged stress. Various forms of social support are also observed in animals under natural conditions (e.g. birds: (Hammers & Brouwer, 2017); pigs: (Cordoni et al., 2023); primates: (Romero et al., 2010)), and experimental paradigms confirm that social support also helps animals to cope with stressful events, through processes referred to as “social buffering” (reviewed in (Kikusui et al., 2006; Wu, 2021)). Social buffering can involve passive processes, whereby the mere presence of a conspecific who is not actively doing anything can help to alleviate stress, or active processes involving direct interactions between non-stressed and stressed individuals. When active social buffering involves affiliative behaviours selectively directed by non-stressed individuals (or observers) towards stressed individuals (or targets), which help to alleviate behavioural and/or physiological signs of stress and fear in targets, such actions fulfil the criteria for “consolation behaviour” (Burkett et al., 2016; de Waal & Preston, 2017).

Active social buffering requires the observer to experience some degree of emotional contagion, defined as an automatic, state-matching response to another’s distress (Adriaense et al., 2020; Düpjan et al., 2020; Palagi et al., 2020; Pérez-Manrique & Gomila, 2018; Pérez-Manrique & Gomila, 2022; Reimert & Bolhuis, 2024). Emotional contagion involves neurophysiological and behavioural changes, and has been documented in many animal species (Adriaense et al., 2020; Pérez-Manrique & Gomila, 2022). Without effective emotional regulation, this contagion can escalate into high-arousal personal distress, leading to self-focused behaviours (e.g., avoidance or general agitation), as the observer attempts to alleviate its own negative state (Clay & de Waal, 2013; Pérez-Manrique & Gomila, 2018). In contrast, consolation requires some degree of emotional regulation to maintain arousal at a moderate level, facilitating other-oriented prosocial responses rather than self-focused ones (Ben-Ami Bartal et al., 2016; Pérez-Manrique & Gomila, 2018). In these cases, the affiliative interaction is initiated spontaneously by the observer, which reduces anxiety levels in the distressed partner (Adriaense et al., 2020; de Waal & Preston, 2017; Pérez-Manrique & Gomila, 2018; Waal, 2008). Evidence for consolation behaviour has been reported in some taxa (voles: (Burkett et al., 2016); primates: (Romero et al., 2010); pigs: (Cordoni et al., 2023)) and it remains of special interest as a potential evolutionary building block for human empathy, bridging emotional reactivity with cognitive perspective-taking (de Waal & Preston, 2017).

The role of complex empathic processes in herd species like farm animals remains poorly understood. Evidence that stable social groups are beneficial for farm animals is based primarily on observed decreases in agonistic behaviours in stable groups compared to groups that undergo remixing (Lee et al., 2022), while effects of group stability on positive affiliative behaviours remains limited. Studies to date show mixed results for evidence of social buffering in different farm animals, but this may also be related to the methods. Most farm animal studies compare subjects who go through stressful events, such as disbudding (calves: (Bučková et al., 2022)), separations from their social group (sheep and goats (Lyons et al., 1993)), or tests in novel environments (chickens: (Bryan Jones & Merry, 1988)) while in pairs versus singly. However, this approach ignores the evidence that individuals who have also undergone the stressor themselves are less effective in providing post-stress social support compared to naïve individuals (Denommé & Mason, 2022).

### 1.2 Social and emotional behaviour in pigs

Domesticated pigs maintain much of the behavioural repertoire of their wild ancestors, and given the opportunity will organize into similar social groupings and exhibit similar social behaviours as wild boars (Gonyou, 2009; Stolba & Wood-Gush, 1989). Among the more common social interactions are affiliative nose-to-nose and nose-to-body contacts, which are used as a greeting and to maintain social cohesion (Camerlink & Turner, 2013). Even brief separations from their social group are stressful for young and growing pigs (Moscovice et al., 2022; Reimert et al., 2014; Ruis et al., 2001), who respond with behavioural indicators of negative affect including escape behaviour and distress vocalizations (Moscovice et al., 2023; Reimert et al., 2013). Brief social separations are also associated with physiological indicators of arousal in pigs, based on rapid increases in glucocorticoid hormones including cortisol, which are released peripherally after activation of the hypothalamic-pituitary-adrenal (HPA)-axis in response to challenges (Herskin & Jensen, 2002; Moscovice et al., 2022).

Research conducted by Reimert and colleagues (Reimert et al., 2013, 2015, 2017) provided foundational evidence that domestic pigs exhibit reliable behavioural and physiological indicators of emotional contagion, including increases in vigilance, ears pinned back and salivary cortisol in naïve pigs after they observed pen mates in negative contexts (Reimert et al., 2013, 2015). These effects also occurred when naïve pigs had not witnessed the stressor directly, but only interacted with pen mates after their return to the home pen (Reimert et al., 2017). These findings established that pigs are sensitive to the negative emotional states of group members, which is a crucial aspect of empathic responses (Pérez-Manrique & Gomila, 2022).

Several studies indicate that the mere presence of social partners leads to social buffering of behavioural (Reimert et al., 2013) or physiological (Kanitz et al., 2014; Tuchscherer et al., 2016) indicators of distress in pigs. To date only one study has tested for evidence for consolation behaviour in domestic pigs. Cordoni and colleagues (Cordoni et al., 2023) demonstrated that in semi-free ranging conditions, third party observers of aggressive interactions sometimes initiate affiliative nose-to-nose or nose-to-body contacts with the losers of these interactions. Losers who receive these interventions have reduced rates of self-directed behaviours that are associated with anxiety, such as head or body shaking, body rubbing and yawning (Cordoni et al., 2023). However, if losers initiate these third-party contacts, they are not effective in reducing anxiety measures. Previous research has also shown that pigs are capable of helping a group member in need, especially when the distressed individual displays clear signs of discomfort [36]. The combined evidence suggests that pigs can perceive the distress of others and potentially regulate their own emotions in order to respond in a goal-directed manner aimed at alleviating others’ distress.

### 1.3 Research Goals

Our main goal was to experimentally test for consolation behaviour in domestic pigs. We aimed to further examine the relationship between emotional contagion, emotional regulation and consolation. To achieve this, we designed a within-subject comparison of observer responses to the same group member after the focal pig was separated and then reunited with their group under two contexts. In one context (direct-reunion), observers remained naïve to the focal pig’s experience before the reunion. In the second context (helping), observers were exposed to various cues that could allow them to assess the emotional state of the focal pig before it was reunited with the group, and they could also control the exact timing of the reunion. In both contexts, we measured changes in affiliative and non-affiliative interactions initiated and received by the reunited individual, compared to matched control observations in an undisturbed social group. We predicted that selective increases in affiliative behaviours by observers without corresponding increases in non-affiliative behaviours would be consistent with moderate, but not high arousal, which is one of the criteria for consolation behaviour (Pérez-Manrique & Gomila, 2018). In contrast, heightened arousal and general social interest in the returning pig were expected to result in a broader spectrum of positive and negative social behaviours (Calderon et al., 2016; Guevara et al., 2024; D. Pfaff, 2009; D. W. Pfaff & Fisher, 2012). We also examined physiological and behavioural indicators of stress reactivity in focal pigs in the two experimental contexts.

We tested whether responses between observer and focal pigs under the different contexts fulfilled three key criteria related to consolation behaviour (Pérez-Manrique & Gomila, 2018): 1) Increases in rates of uni-directional affiliative behaviour from observers towards the distressed individual (but without corresponding increases in affiliation from distressed individuals to observers); 2) No changes in rates of non-affiliative interactions from observers to distressed individuals, and 3) Reduced anxiety behaviours and reduced cortisol reactivity in focal pigs who receive affiliative interactions (see Table 1). We hypothesized that evidence for consolation behaviour would be stronger in the helping context, where additional cues were available to assess the focal’s emotional state and observers had more control over when they interact with the separated pigs. Both of these aspects may promote the emotional regulation required for consolation behaviours. Alternatively, it is possible that greater exposure to distress cues from others, without the ability to regulate emotions, may increase personal distress and impede prosocial responses, as observed in previous studies (Ben-Ami Bartal et al., 2016). (insert Table 1)

**Table 1.** Criteria evaluated for consolation in pigs^1^, and comparison of evidence in support of the criteria across two reunion contexts: Direct-reunion (without prior social cues available to observers) vs. Helping contexts (with prior social cues available to observers). A check mark (✓) indicates that the data support the hypothesis, a cross (✗) indicates a lack of support.

| Criterion | Predicted outcome | Direct-Reunion outcome | Helping outcome |
| --- | --- | --- | --- |
| 1. Uni-directional affiliation toward the distressed individual (focal) | 1a. Increase in affiliative contacts received by focal | ✓ | ✓ |
|  | 1b. No increase in affiliative contacts given by focal | X | ✓ |
| 2. Moderate arousal in bystanders (observers) | No increase in non-affiliative contacts received by focal | X | ✓ |
| 3. Stress alleviation in the distressed individual (focal) | 3a. Focals show reduced post-reunion anxiety behaviours | X | ✓ |
|  | 3b. Focals show reduced post-reunion cortisol concentrations | X | X |

<sup>1</sup>Criteria are based on the framework proposed by Pérez-Manrique and Gomila 2018

## 2. Methods

### 2.1 Animals and husbandry

The research was conducted in accordance with the German Animal Welfare Act (German Animal Protection Law, §8 TierSchG) and was approved by the Committee for Animal Use and Care of the Agricultural Department of Mecklenburg-Western Pomerania, Germany (permits: LALLF 7221.3-1-036/20). Subjects were German Landrace pigs born and housed at the Experimental Pig Facility at the Research Institute for Farm Animal Biology (FBN) in Dummerstorf, Germany. Husbandry conditions adhere to the standards for conventional housing of pigs in Germany. After birth, piglets are housed with their mothers in farrowing compartments (6 m^2^) with plastic floors and a covered, heated piglet nest area (0.90 m^2^). Piglets have unrestricted access to water and are offered dry food (HAKRA-Immuno-G; UnaHakra, Hamburg, Germany) in addition to milk beginning from post-natal day (PND) 14. Piglets receive tattoos on PND 1 and ear tags with identification numbers at PND 27. Pigs are not subjected to tail docking, teeth clipping or castration. At PND 28, piglets were weaned and mixed into new social groups, by randomly selecting either three pigs from three different litters, or five pigs from two different litters, resulting in groups of 9-10. The choice of mixing pigs from two or three different litters was based on the availability of litters with the appropriate number of piglets to maintain sex-balanced groups, in which each pig had equal numbers of littermates present. Groups were sex balanced and contained equal numbers of littermates from each contributing litter, such that each pig had either two (22% of group members) or four (44% of group members) relatives in their group. For the current study, we analysed additional data from two previously published experiments (Moscovice et al., 2022, 2023). This new analysis combines the datasets and applies a novel perspective to address a different research question. In doing so, we embrace the principle of Reduction, one of the foundational elements of the 3Rs, which guide ethical animal research (Russell & Burch, 1959). Re-examining existing data through new conceptual lenses allows us to advance knowledge while minimizing the ethical and practical costs associated with animal experimentation.

### 2.2 Experimental procedures

The study design maintains experimental control while preserving typical group-living conditions in commercially farmed pigs. Pigs were tested in four temporally separated cohorts, based on the five-week interval between the births of new litters. Each cohort consisted of two social groups, with 9-10 pigs per group, for a total of n = 78 subjects. One pig died of unknown causes, and two pigs from the same group were removed from the study to be treated for illnesses before testing began. Due to technical problems, behavioural data from one pig was not recorded during the isolation experiment. We therefore report the results from the remaining n = 74 pigs who completed both experiments. Groups within a cohort were housed in neighbouring 2.8 m^2^ pens in the same room, with partial slatted flooring and a sleeping mat. Pigs could receive auditory and olfactory cues from neighbouring pens, but did not have direct visual or physical contact with members of the other pen. Pigs were regularly given back numbers using livestock spray paint to facilitate individual identification. The room was equipped with multiple video cameras to cover the entire home pen and two additional compartments that were later attached to the pen. Each experiment was conducted across five consecutive days, between 9-12 hr. On each test day, both groups within a cohort were tested in a counter-balanced order, with two subjects per group tested as the focal pigs. Each focal pig was tested once in each experiment, and once on a time-matched control day, in a within-subject design (**insert Figure 1**).

**Figure 1.**
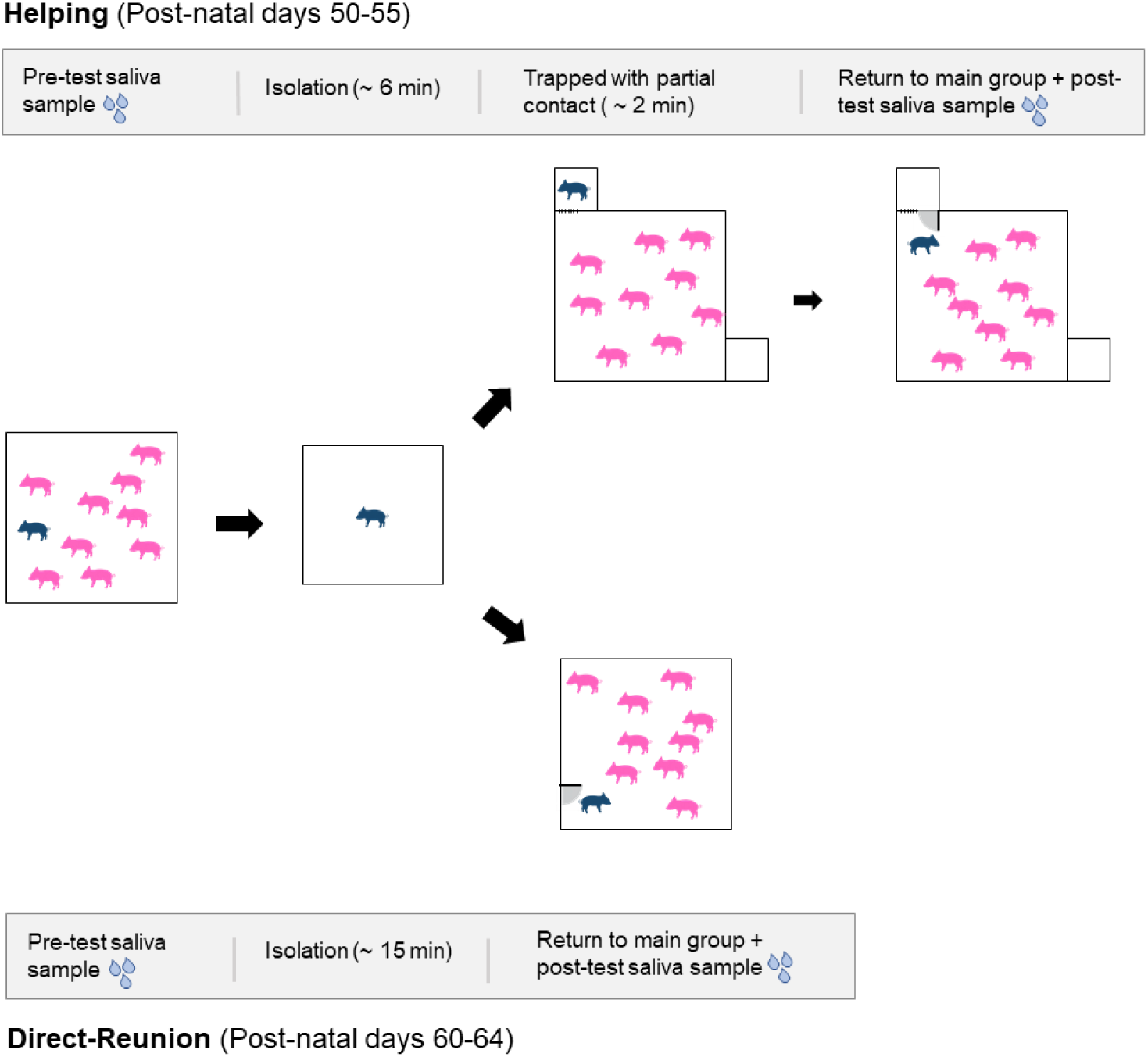
Comparison of testing procedure for focal and observer pigs in the Helping and Direct-reunion contexts

#### 2.2.1 Control context

We conducted one 5-min behavioural observation on each focal subject using pre-recorded video from the home pen. Observations occurred between 45–49 days of age, while all pigs were in their stable social groups and before any social separation events occurred. We used different days and times for each focal pig, and matched the timing of the video observation during the day with the timing of each pig’s direct-reunion test (see below).

#### 2.2.2 Helping context

Between 50 and 55 days of age, pigs were tested in a prosocial task (Moscovice et al., 2023). Prior to this, pigs had been habituated to interactions with one of the researchers (LRM) and one assistant, who interacted with the pigs daily for two weeks. Pigs also had one week of exposure in their social groups to two identical, novel compartments that were attached to different sides of the home pen. Each compartment had a mesh window, and a door with a metal handle, that when lifted high enough released an inner latch, causing the door to the compartment to swing open into the home pen. During their one-week exposure, pigs could learn a novel behavior to open the compartment doors and explore inside. On each day of the following test week, one of the researchers (LRM) or an assistant entered the stall and collected a saliva sample from the pre-determined focal pigs within the group. Pigs had been previously habituated to this procedure and voluntarily chewed on swabs to provide saliva samples within 2 min without requiring restraint (Moscovice et al., 2022). After collecting the saliva samples, one of the two familiar experimenters entered the stall, picked up the first focal pig, carried the pig out of the stall and room, and placed them in a novel room, in a 2.8 m^2^ empty arena. Pigs remained isolated in the room for median = 5.8 (IQR = 5.75–6.06) min. The experimenter then entered the room, picked up the pig and carried it back to the home pen room, and placed the pig in one of the two compartments. From the compartment, the focal pig could maintain visual and limited physical contact with group members through the mesh window. Pigs remained trapped in the compartment for median = 2.2 (IQR = 1.27–4.66) min. The trial ended when one of the group members opened the compartment door, releasing the focal pig back to the group. If focal pigs were not released within 20 min, a researcher (LRM) or assistant opened the compartment door to release the pig back to the group. Once the door of the test compartment was opened, pigs were left undisturbed for an additional 15 min reunion period and then a post-experiment saliva sample was collected, representing cortisol reactivity approximately 20–40 min following the onset of the isolation stressor. Two focal pigs per group were tested each day, with an approximately 1 h pause between the first test and the second test within a group. Testing continued for five consecutive days, until every pig in the group had been isolated and reunited once.

#### 2.2.3 Direct-Reunion context

Between 60 and 64 days of age (five days following the completion of the helping experiment), each pig underwent one 15-min separation from their social group in a pre-determined random order (Moscovice et al., 2022). Before the separation, a researcher (LRM) or assistant entered the pen and collected a baseline saliva sample from the focal pig, as described above. The person then opened the door to the home pen and allowed the focal pig to leave the pen, which pigs did willingly without being handled. A sorting board was used to encourage the pig to walk down a short hallway until they entered an open door to the isolation room, containing the empty arena where the pigs had been isolated during the helping experiment. The arena door was closed and the focal pig was left alone for 15 min. The person then returned to the room, collected a post-isolation saliva sample and led the pig down the hallway and back into their home pen. Pigs voluntarily walked down the hallway and returned to their home pen without being carried. Pigs were left undisturbed for the 15-min reunion period, after which a post-reunion saliva sample was collected from the focal pig, representing cortisol reactivity approximately 30 min following the onset of the isolation stressor. We tested two subjects per group per day and alternated between groups, so that each group had a recovery period of ∼ 45 min between the end of a reunion and the next isolation. Testing continued for five days, until each pig had been isolated and reunited once.

### 2.3 Behavioural data

Behavioural data were coded from videos using focal continuous sampling methods and the Observer software (version 15, Noldus, The Netherlands). For experiments, coding began at the moment when the door from the compartment with the trapped pig was opened (helping) or immediately when the focal pig’s head had entered the home pen (direct-reunion). Each focal animal was observed for 5 min.

For the behavioural data, we followed the ethogram of Cordoni et al. (Cordoni et al., 2023) with some modifications due to the more limited space constraints in our captive groups. We focused on behaviours across three categories: affiliative interactions, non-affiliative interactions (including ambiguous and aggressive interactions) and anxiety behaviours (for complete Ethogram, see Supplementary Table S1). For social interactions, we recorded the identity of the partner and the direction of the interaction, based on who initiated the contact. Thus, for example for nose-body contact, the initiator was defined as the first pig in the dyad to turn its head towards its partner and make contact with the body. In the case of non-affiliative or aggressive behaviour, it was the pig that initiated the interaction either by chasing, pushing or directing its head towards the opponent first. Similarly to Goumon et al. (Goumon et al., 2020), we did not assess directionality for nose-nose contacts, since in at least half of the cases it was not possible to accurately identify the initiator of the contact. Additionally, vocalizations were not coded because the group setting prevented reliable attribution of calls to the focal pig.

Videos were coded by one of the authors (WJ) and inter-observer reliability was checked by another author (ALC), which resulted in a high agreement score (kappa = 0.84). Due to differences in the testing procedures, coders could not be blind to the different contexts. All behaviours included in the analysis were coded as point events. Two occurrences of the same behaviour involving the focal pig were coded as two independent events if the interval between them was at least 5 s, or if they involved a new social target (for social behaviour). For the purposes of the analysis, we summed behaviours within each behavioural category, resulting in a database including for each focal, their total number of affiliative interactions (including directional and non-directional affiliation), total number of directional affiliative interactions given and received (excluding nose-nose contacts), total number of non-affiliative/aggressive behaviours given and received and total number of anxiety behaviours. We also recorded the latency to first affiliative contact for each focal. We present analyses of the directional affiliation data, which is most relevant for evaluating our predictions about consolation behaviour. However, models testing for effects of context on total affiliative behaviour (including non-directional nose-nose contacts) showed similar patterns (see Supplementary Tables S2 and S3).

### 2.4 Dominance ranking

On PND 29–30, during the first two full days following weaning, two assistants recorded all occurrences of fights (as defined in the ethogram) from video data with a high agreement score (kappa = 0.79). Fights were recorded for two hours per day, divided between morning and afternoon observations, resulting in four observation hours per group. During that time, n = 478 fights (group median = 48.5, IQR= 42.5–64.5) were recorded. For decided aggressions, the winner was identified as the pig who eventually displaced or chased away another. Undecided outcomes, in which neither opponent was clearly displaced or showed submission, were recorded as ties. Dominance was assessed using the EloRating package (Neumann et al., 2011), with outcomes scaled between 0 and 1.

### 2.5 Hormone analyses

Saliva samples were collected voluntarily from pigs using SalivaBio® Infant Swabs (Salimetrics, CA, USA). Cortisol analyses were conducted at the FBN, using an ELISA kit for salivary cortisol (Demeditec Diagnostics®, GmbH, Kiel, Germany) that has been previously validated by our laboratory in pigs (Goursot et al., 2019; Moscovice et al., 2022). The standard curve ranges from 0.1–30 ng ml^−1^ and assay sensitivity is 0.019 ng ml^−1^. We compared changes in cortisol in samples collected from each focal pig during the helping and direct-reunion experiments. Changes were based on differences from pre-test samples, taken before the experiment began and post-reunion samples, taken 15 min after the focal pig had full access to their home pen again. In each context, post-test samples occurred between 20-40 min following the onset of the isolation stressor, which is within the reported time interval for peak salivary cortisol responses to stressors in humans (Wirobski et al., 2024)). We have previously reported detailed methods and validations of the cortisol analyses in pigs (Goursot et al., 2019; Moscovice et al., 2022). In brief, samples were thawed, spun at 2,500 × g for 5 min and 50 μl of supernatant was diluted 1:2 with assay buffer. Samples were then run in duplicate. The inter-assay CVs of low- and high-concentration pooled controls were 9.3% and 8.3%, respectively (n = 16 samples). Intra-assay CVs of low- and high-concentration pooled samples were 10.7% and 8.6%, respectively.

### 2.6 Statistical analysis

To test for influences on directional affiliation, latency to first directional affiliative contact (log-transformed) was modelled with a linear mixed model (LMM) containing experimental context × direction of affiliation, sex and rank. Counts of affiliative interactions initiated and received were analysed using a zero-inflated negative binomial generalized linear mixed model (GLMM) with experimental context × direction of affiliation, kinship, sex, and rank as fixed effects. To control for varying availability of potential kin partners, we included the log-transformed count of number of kin present as an offset term. To control for general arousal of observer pigs, influences on directional non-affiliative interactions were modelled using the same structure. Because anxiety behaviours were rare, they were dichotomized (present/absent) and analysed with a binomial GLMM that included affiliation received × context, non-affiliative behaviour received × context, sex, and rank as fixed effects. Salivary cortisol concentrations were log-transformed. Due to missing samples from three pigs and an outlier measure, the sample size for these models was reduced to n = 280 samples from n = 70 pigs (two samples per pig per context). The initial LMM included sample type (pre- vs post-experiment) × rate affiliation received, sample type × rate non-affiliative behaviour received, sample type × anxiety (present/absent) and sample type × context (helping vs direct-reunion). We also included sample type × time interval between initial separation stressor and saliva sampling, to control for any effects of differences in duration of the stress events on measures of cortisol concentrations. Sex, rank and time of day of sample collection were included as additional control predictors.

In all models, focal pig ID and group were entered as random effects unless their inclusion caused singularity issues. In such cases, the factor with lower variance was removed. All behavioural variables were first inspected graphically. Skewed responses were normalised (square-root or log), and continuous predictors (rank and sampling time in the cortisol analysis) were z-transformed (mean = 0, SD = 1) to aid convergence and interpretation. Results are reported as medians and inter-quartile ranges (IQRs) unless otherwise stated. Model assumptions were checked with residual plots, Q–Q plots, over-dispersion statistics, and variance-inflation factors (VIF < 2). Estimated marginal means (EMMs) and pairwise contrasts were computed with the emmeans (Lenth et al., 2025) package using Tukey’s adjustment for multiple comparisons. For all models we report fixed-effect coefficients and p-values. All statistical procedures were carried out in R (v 4.3.2) (*R: The R Project for Statistical Computing*, n.d.). Data handling used tidyverse (Wickham et al., 2019), mixed-effects modelling used lme4 (Bates et al., 2015), lmerTest (Kuznetsova et al., 2017), and glmmTMB (Brooks et al., 2017); diagnostic checks used performance (Lüdecke et al., 2021) and multicollinearity was assessed with car (Fox et al., 2024). All data and scripts related to analyses are available at: https://doi.org/10.5281/zenodo.18336668

## 3. Results

Of the focal (separated) pigs, 81% (n = 59) were involved in affiliative interactions (given or received) during control observations, while these numbers increased to 97% (n = 72) following helping and 100% (n = 74) following direct-reunions. For the focal pigs who were involved in affiliative interactions during observations, latency to affiliate tended to be influenced by an interaction between context and direction of affiliation (LMM, F_2,353_ = 2.407, p = 0.092, see Supplementary Table S4). Focal pigs had shorter latencies to receive affiliation during helping (27.4 (8.0–58.1) sec) or direct-reunions (10.4 (4.8–27) sec) compared to control contexts (67.6 (17.8–154) sec, t ratios > 4.02, adj. p values < 0.001). Focal pigs also had shorter latencies to receive affiliation following direct-reunions compared to helping contexts (t-ratio = 3.02, p = 0.007). Focal pigs were also faster to initiate affiliation following direct-reunions compared to helping and to control contexts (t-ratios > 5.81, adj. p values < 0.001). However, focals did not differ in latencies to initiate affiliation between helping and control contexts (t-ratio = 1.74, p = 0.188, see Supplementary Table S5).

There was a significant interaction of context and direction of interactions on the amount of directional affiliation that focal pigs were involved in (GLMM, χ^2^ = 16.967, p < 0.001, see Table 2 and Figure 2). Post hoc tests indicate that focal pigs received more directional affiliation during both helping (Median = 1 (IQR = 0.4–1.6) interactions/min) and direct-reunions (1.8 (1–2.4) interactions/min) compared to control contexts (0.4 (0–0.8) interactions/min, z-ratios < −6.7, adj. p values < 0.001, see Supplementary Table S6). They also received more directional affiliation during direct-reunions compared to during helping (z-ratio = −4.1, adj. p < 0.001). However, focal pigs also initiated more affiliation to other pigs during direct-reunions (1.2 (0.8–1.6) interactions/min) than during helping (0.4 (0–0.8) interactions/min) or control (0.2 (0–0.8) interactions/min) contexts (z-ratios < −7.85, adj. p values < 0.001). In contrast, there were no differences in amount of directional affiliation given by focal pigs to other group members (observers) between helping and control contexts (z-ratio = −0.92, adj. p = 0.627, see Supplementary Table S6). (insert Table 2 and Figure 2)

**Figure 2.**
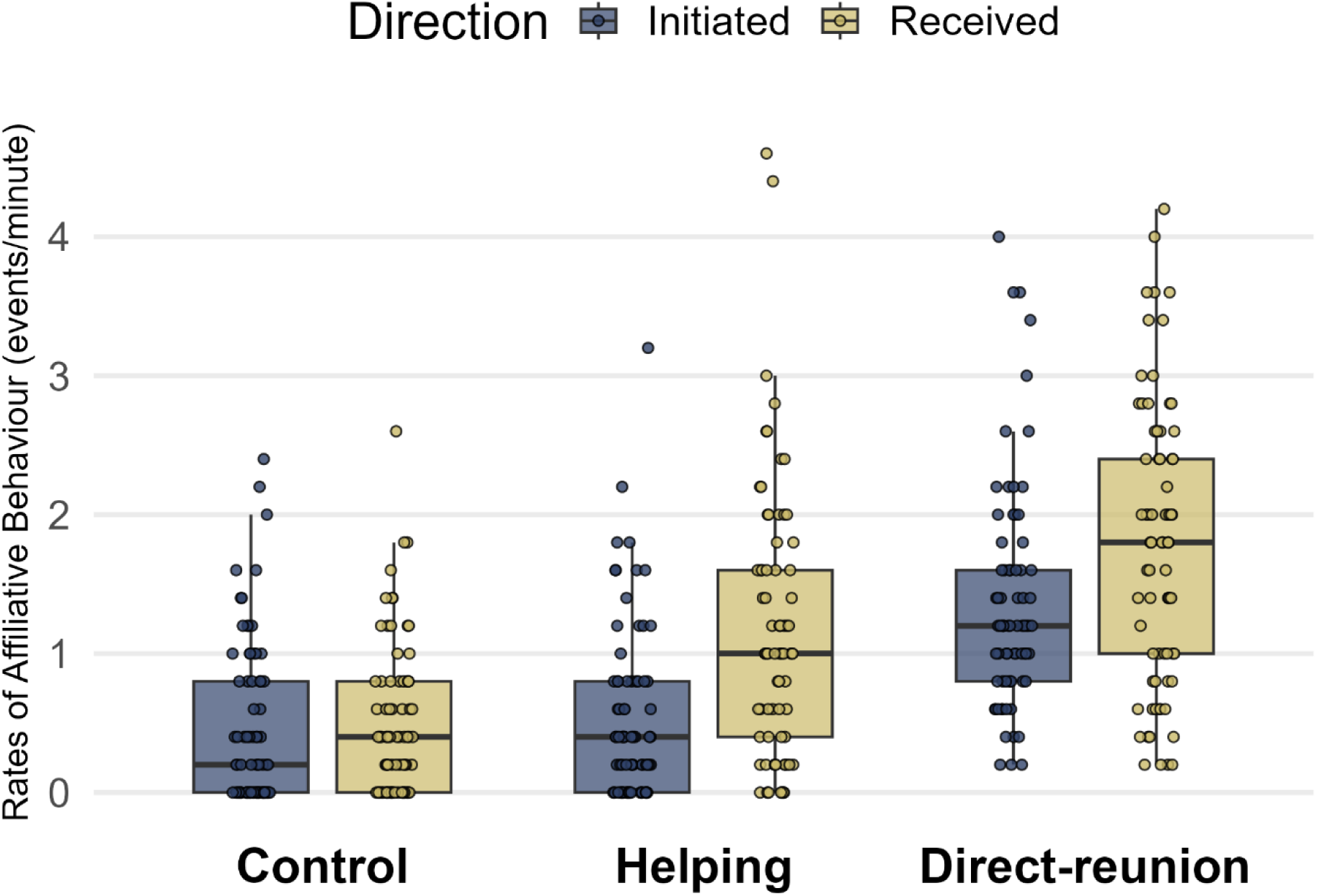
Rates of directional affiliative behaviour given and received by the focal pigs (n = 74) across the control, helping and direct-reunion contexts. See also Table 2.

**Table 2.** Results of a GLMM testing for an interaction between context (control, helping, direct-reunion) and direction of affiliation (given or received) on amount of affiliation between focal pigs and their group members. Significant predictors are indicated in bold.

| predictor | estimate | s.e. | z or $\chi^2$<br>value | *p*-value |
| --- | --- | --- | --- | --- |
| (Intercept) | 0.915 | 0.119 |  | (a) |
| context_helping | 0.823 | 0.1518 |  | (a) |
| context_direct-reunion | 1.251 | 0.122 |  | (a) |
| direction_give | -0.061 | 0.135 |  | (a) |
| sex_female | -0.052 | 0.083 | -0.632 | 0.527 |
| elo rank | -0.020 | 0.042 | -0.468 | 0.64 |
| kinship_yes | 0.033 | 0.067 | 0.486 | 0.627 |
| **context_helping:direction_give** | -0.700 | 0.181 | 16.967 | **< 0.001** |
| **context_direct-reunion:direction_give** | -0.198 | 0.169 |  | (b) |

<sup>a</sup> Not indicated due to limited interpretability, either of intercepts or due to significant interaction terms
<sup>b</sup> Refer to $\chi^2$ value for 'context\_helping:direction\_give' for overall effect of interaction

Non-affiliative interactions consisted primarily of low-intensity ambiguous or aggressive behaviours, with only two instances of fighting recorded. There was no effect of context on the amount of non-affiliative interactions given or received for focal pigs (GLMM, context*direction, Likelihood ratio test, χ^2^_(2)_ = 0.688, p = 0.702, see Figure 3). With the non-significant interaction removed, there was a main effect of context on overall non-affiliative interactions (GLMM, χ^2^_(2)_ = 15.578, p < 0.001, see Table 3). Post-hoc tests indicate that focal pigs were involved in more non-affiliative interactions (given and received) following direct-reunions (0.2 (0.2–0.75) interactions/min), compared to helping (0.2 (0–0.4) interactions/min), z-ratio = 2.39, adj. p = 0.044) and to control observations (0 (0–0.2) interactions/min), z-ratio = 3.87, adj. p < 0.001). Rates of non-affiliative interactions did not differ between helping and control observations (z-ratio = −1.55, adj. p = 0.266, see Supplementary Table S7). In addition, focal females tended to be involved in fewer non-affiliative interactions than males (GLMM, est ± se = −0.29 ± 0.17, p = 0.098), which is consistent with previous literature (Colson et al., 2006). (insert Table 3 and Figure 3)

**Figure 3.**
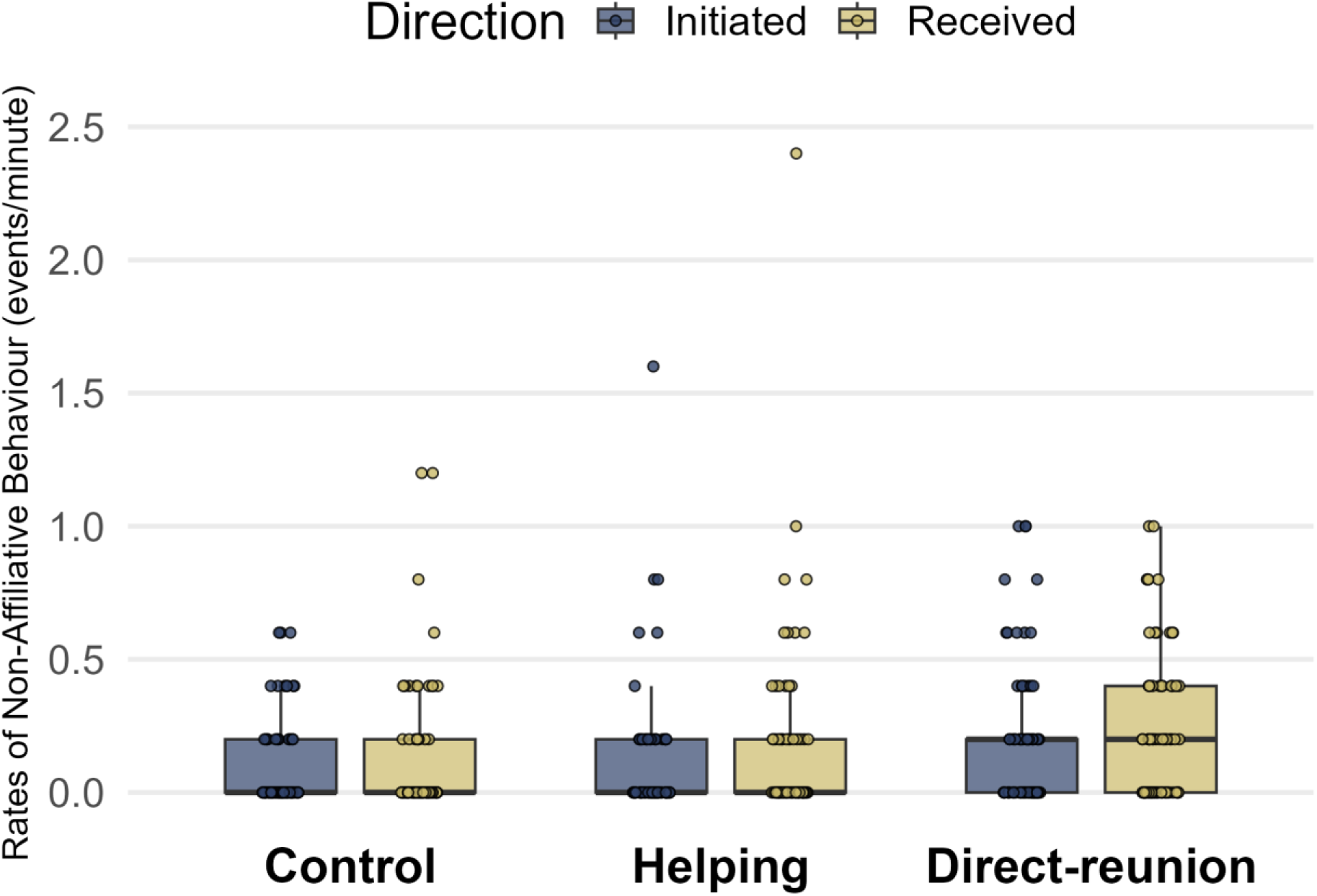
Rates of directional non-affiliative behaviour given and received by the focal pigs (n = 74) across the control, helping and direct-reunion contexts. See also Table 3.

**Table 3.**
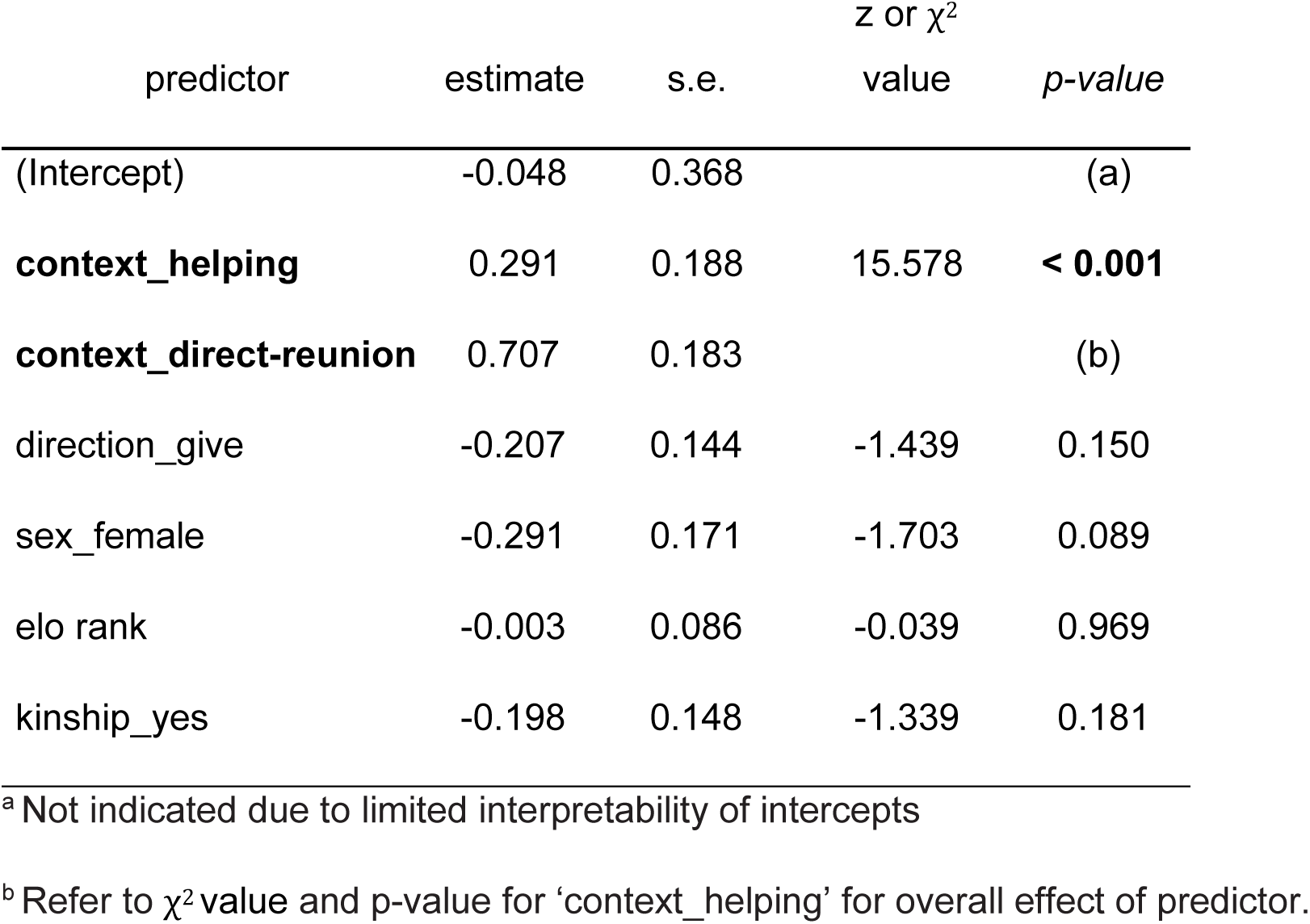
Results of a GLMM testing for main effects of context (control, helping, direct-reunion) and direction of interactions (given or received) on amount of non-affiliative interactions between focal pigs and their group members. Interaction terms were not significant and were removed from the final model. Significant predictors are indicated in bold.

| predictor | estimate | s.e. | z or $\chi^2$<br>value | *p*-value |
| --- | --- | --- | --- | --- |
| (Intercept) | -0.048 | 0.368 |  | (a) |
| **context_helping** | 0.291 | 0.188 | 15.578 | **< 0.001** |
| **context_direct-reunion** | 0.707 | 0.183 |  | (b) |
| direction_give | -0.207 | 0.144 | -1.439 | 0.150 |
| sex_female | -0.291 | 0.171 | -1.703 | 0.089 |
| elo rank | -0.003 | 0.086 | -0.039 | 0.969 |
| kinship_yes | -0.198 | 0.148 | -1.339 | 0.181 |

<sup>a</sup> Not indicated due to limited interpretability of intercepts
<sup>b</sup> Refer to $\chi^2$ value and *p*-value for 'context\_helping' for overall effect of predictor.

Across the different contexts, we recorded n = 91 anxiety behaviours, the majority of which involved body shaking (74%), followed by body scratching (19%). Yawning (n = 5 occurrences) and vacuum chewing (n = 2 occurrences) occurred rarely, and with one exception were observed only in control contexts. In models explaining anxiety behaviours in focal pigs, after removing non-significant interactions between context and rates of affiliation received (GLMM, Likelihood ratio test, χ^2^ = 4.154, p = 0.125) and context and rates of non-affiliative behaviour received (GLMM, Likelihood ratio test, χ^2^ = 0.916, p = 0.632), the final model indicated that anxiety was influenced by a main effect of context (GLMM, χ^2^ = 7.100, p = 0.029, see Table 4), due to more focal pigs showing anxiety behaviours following direct reunions (39% of pigs), compared to following helping (19% of pigs, z-ratio = 2.53, adj. p = 0.031, see Figure 4 and Supplementary Table S8). However, there were no significant differences in the likelihood of anxiety behaviours between the two tests and control observations, in which 28% of pigs showed anxiety behaviours (z-values < 1.23, adj. p values > 0.422). There was also a main effect of non-affiliative behaviour received, such that focal pigs who received more non-affiliative behaviour across contexts were also more likely to show anxiety behaviours (GLMM, est ± se = 0.57 ± 0.17, p = 0.001, see Table 4). (insert Table 4 and Figure 4)

**Figure 4.**
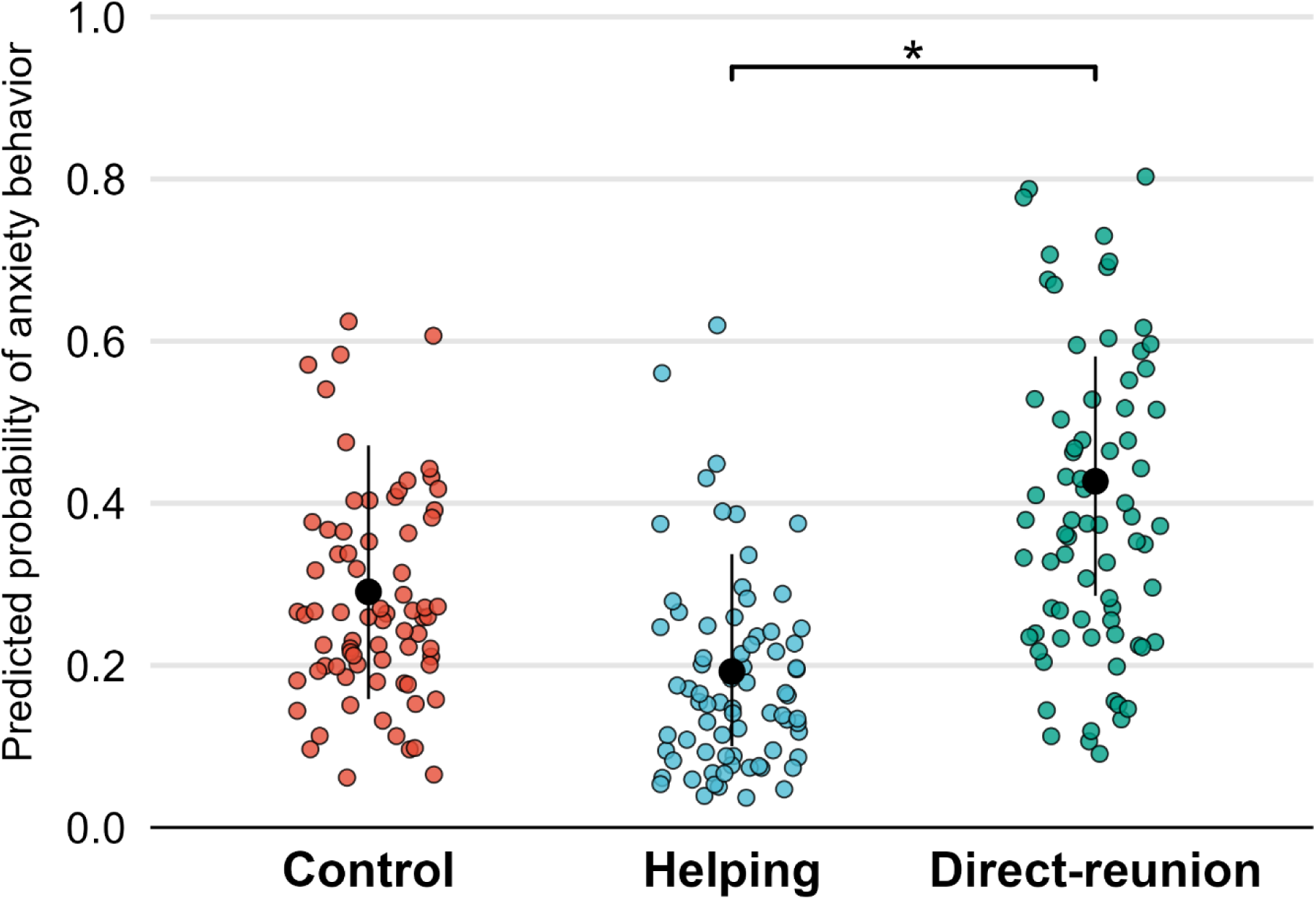
Likelihood of anxiety behaviors across contexts. Individual predicted probabilities (colored dots) and estimated marginal means with 95% confidence intervals (black point-ranges) are derived from the GLMM analysis (see Table 4). Asterisks indicate significant differences based on post-hoc comparisons (see Supplementary Table S8).

**Table 4.** Results of a GLMM testing for interactions between context (control, helping, direct-reunion) and affiliative or non-affiliative social interactions received on expression of anxiety behaviours in focal pigs. Significant predictors are indicated in bold.

| predictor | estimate | s.e. | z or $\chi^2$ | |
| --- | --- | --- | --- | --- |
|  |  |  | value | *p value* |
| (Intercept) | -1.290 | 0.421 |  | (a) |
| affiliation received | -0.274 | 0.218 | -1.253 | 0.200 |
| **non-affiliative received** | 0.565 | 0.174 | 3.243 | **0.001** |
| **context_helping** | -0.542 | 0.441 | 7.100 | **0.029** |
| **context_direct-reunion** | 0.598 | 0.478 |  | (b) |
| sex_female | 0.383 | 0.350 | 1.092 | 0.274 |
| elo rank | -0.297 | 0.183 | -1.624 | 0.097 |

<sup>a</sup> Not indicated due to limited interpretability of intercepts
<sup>b</sup> Refer to $\chi^2$ value and p-value for 'context\_helping' for overall effect of predictor.

In both helping and direct-reunions, focal pigs exhibited increases in salivary cortisol in samples taken between 20–40 minutes following the onset of the isolation stressor (see Supplementary Fig. S1). However, in models predicting cortisol reactivity, there were no significant interactions between rates of affiliation received, rates of non-affiliative interactions received, test context (direct-reunion vs. helping) or time interval between onset of the separation stressor and the sampling time on changes in cortisol from pre- to post-test (LMM, χ^2^ values < 2.51, p values > 0.113). After removing non-significant interactions, in the final model, pigs who showed anxiety behaviours differed in their cortisol reactivity from pigs who did not show anxiety behaviours (LMM, est ± se = 0.22 ± 0.1, p = 0.025, see Supplementary Table S9). Post-hoc tests revealed that the significant interaction was due to pigs showing anxiety behaviours having higher pre-test cortisol concentrations (t ratio = −3.123, p = 0.002, see Supplementary Table S10). There were no other effects of predictors on changes in cortisol, but total cortisol concentrations tended to be higher in focal pigs who received more non-affiliative behaviour from group members (LMM, est ± se = 0.06 ± 0.03, p = 0.051, see Supplementary Table S9).

## 4. Discussion

This study reveals context-dependent modulation of affiliative as well as non-affiliative interactions in domestic pigs, providing insight into the circumstances under which responses may (and may not) qualify as consolation behaviour. The helping context produced a selective increase in affiliation directed toward the distressed focal pig without corresponding increases in non-affiliative contacts and was associated with lower anxiety behaviours in focal pigs, when compared with the direct-reunion context (see Supplementary video). This evidence aligns with the pattern expected in consolation (see Table 1), whereas results in the direct-reunion context may be better explained by emotional contagion leading to high levels of arousal and personal distress.

Focal pigs were quicker to receive affiliation and received the most affiliative contacts after a direct-reunion, followed by the helping context, and least after the control context. This gradient appears to contradict our predictions that helping contexts, during which observer pigs received more cues about the emotional state of pigs in need of help, would facilitate consolation behaviour. However, direct-reunions also led to significant increases in affiliation given by the focal pig to other group members, as well as in non-affiliative interactions given and received, compared with the helping context and the control observations. This combined evidence is consistent with emotional contagion leading to high-arousal and personal distress in observer pigs, rather than consolation (Clay & de Waal, 2013; Pérez-Manrique & Gomila, 2018). This interpretation aligns with previous findings in pigs that negative emotional experiences can spread to naïve individuals, evidenced by both behavioural and physiological indicators (Reimert et al., 2013, 2015). Experimental investigations of consolation behaviour often measure the likelihood of affiliative contacts between näive observers and stressed targets following a test vs. control situation, but without examining the occurrence of other behavioural interactions. As a result, such designs may inadvertently label as “consolation”, increases more generally in social interactions that may naturally follow socially-arousing contexts, such as disruptions to the social group or aggressive interactions.

By contrast, the helping context fulfils most of the prerequisites for genuine consolation (see Table 1). First, there was a selective increase in affiliation toward the distressed focal pig soon after reunion, without a matched increase in affiliation from the focal, precluding the possibility that the behaviour was simply a response to solicitation (Pérez-Manrique & Gomila, 2018). If observers were merely over-aroused, the optimal strategy would be to seek affiliation with some of the other calm, unstressed group members present, who would be more effective in social buffering (Kikusui et al., 2006; Wu, 2021), rather than interacting with the distressed focal. The increases in affiliation by observers towards the focal pig, combined with the relatively lower cortisol levels reported in the same observer pigs soon after helping (Moscovice et al., 2023), suggest that observer pigs were not initiating affiliation in order to regulate their own arousal. Second, non-affiliative interactions remained at similar levels as during control days, indicating that observers were not simply more aroused (Wascher, 2021) or more curious about the returning pig. Third, focal pigs showed lower levels of post-reunion anxiety in helping contexts than in the direct-reunion contexts, although we did not find a direct relationship between increased affiliation from observers and reduced anxiety or cortisol in focal pigs, as we had predicted. Overall, the evidence from the helping context is consistent with the idea that some observer pigs perceived the situation of the stressed focal, showed evidence for emotional regulation and reacted in ways appropriate to alleviating distress in the focal pig. They did this by opening a door to release them back to the group (Moscovice et al., 2023) and selectively increasing post-reunion affiliative behaviour towards reunited pigs, who showed little evidence for anxiety after their reunion.

During direct-reunions, focal pigs willingly left and re-entered the home pen on their own and only exhibited clear distress behaviours while isolated in a separate room, which occurred for a set duration of 15 minutes per pig. In the helping context, after the 5.8 min median isolation time in another room, focal pigs were carried back to their home pen room and then spent a variable period of time (median 2.2 min, IQR = 1.27–4.66 min) in a compartment attached to the home pen, where they could maintain visual, auditory and limited tactile contact with their group members before they were reunited with their group. While being carried in and out of the home pen room, and while trapped in the compartment, the majority of focal pigs showed evidence of distress, indicated by squeal and scream vocalizations and escape attempts (Moscovice et al., 2023), all of which were available as cues for observers to assess the emotional state of focal pigs. These additional cues about the emotional state of focals, along with the intervening period in the compartment, likely served several functions. First, the mere presence of group members while focal pigs were trapped in the compartment is likely to have relieved some of the focal pigs’ stress via social buffering (Wu, 2021). Additionally, this period gave observers more opportunities to recognize their returning group member, to assess the trapped pig’s emotional state and to emotionally regulate their own arousal before physical reunion, which may have facilitated targeted affiliative responses (Adriaense et al., 2020; Pérez-Manrique & Gomila, 2018). The combination of these factors may explain why the helping context selectively heightened affiliation received by the focal (but not affiliation given or non-affiliative interactions), and was associated with reduced anxiety in focal pigs, whereas in direct-reunions all measured behaviours were significantly higher compared to the helping context.

Anxiety behaviours are established indicators of stress in pigs (Norscia, Collarini, et al., 2021), and their association with elevated baseline salivary cortisol concentrations in our study further supports this interpretation. The relatively high percentage of focal pigs showing anxiety behaviours in the control context was a surprising result that may be due in part to yawning and vacuum chewing being found almost exclusively in this context; both behaviours have been linked to boredom (Zhitnitskiy et al., 2021) or to emotional communication in pigs (Norscia, Coco, et al., 2021). Across both test contexts, anxious focal pigs received more non-affiliative interactions, but did not receive more affiliation compared to pigs who did not exhibit anxiety behaviours. Moreover, higher overall cortisol concentrations in focal pigs predicted their non-affiliative interactions received. Two non-exclusive mechanisms may account for this pattern. First, elevated cortisol and visible anxiety in the focal pig could, through emotional contagion, lead to high levels of arousal and personal distress in observers, which could increase the likelihood of investigatory or aggressive behaviours toward the focal pig, as well as towards other group-mates (Fernandez et al., 1994; Guevara et al., 2024). Second, pigs subjected to non-affiliative behaviours may themselves perform more anxiety-related behaviours and display elevated cortisol (Otten et al., 1999). Disentangling causality would require more prolonged cortisol sampling of focal pigs beyond 15 minutes post-reunion, to better capture the longer-term effects of different types of social interactions on stress responses. More detailed data on cortisol reactivity and social interactions among the observers themselves would additionally help to determine the extent to which behavioural responses to focal pigs are indicators of more generalized arousal.

Future work should further investigate the relative contributions of (i) the focal pig’s diminishing stress during the social buffering phase next to the pen in the helping context and (ii) the observers’ reaction to that stress, to the initiation of consolation behaviour. During this period, continuous recordings of heart-rate variability (HRV) could offer a second-by-second window onto autonomic regulation in both focal pigs and their group-mates (Krause et al., 2017; von Borell et al., 2007). When these physiological traces are synchronized with automated movement tracking of the focal and group-mates, total locomotor activity could become another quantitative proxy to help explain how much of the social buffering (for focal pigs) and emotional contagion (for observers) in the helping context accrues while the focal pig remains confined to the side compartment (Oczak et al., 2022). Lastly, to enhance causal inference of the role of emotional contagion in consolation, the flow of information between partners could be manipulated directly. We propose contrasting the present two-way compartment, which allows bidirectional visual and olfactory access, with a unidirectional variant fitted with a one-way mirror, tinted on the focal pig’s side, so that pen-mates can still appraise the focal pigs’ state while the isolated animal does not benefit from social buffering. If consolation is driven primarily by the perception of distress in others, then observers should show stronger evidence for consolation behaviour following this one-way mirror condition compared to the two-way condition, where targets can benefit from passive social buffering processes before reunions; if however emotional contagion of distress impedes consolation behaviour, then observers should show weaker evidence for consolation in this one-way condition compared to the two-way condition, due to heightened and prolonged distress in trapped pigs in the one-way mirror condition.

From a husbandry standpoint, our findings emphasize the need to preserve group stability and to handle temporary separations with care. The increase in non-affiliative behaviours observed after the sudden re-entry in direct-reunions has some similarities with increases in aggressive behaviour in commercial farm settings following reunions of group members after longer separations (Ewbank & Meese, 1971). Limiting remixing events across the production cycle reduces stress and aggression in domestic pigs (Coutellier et al., 2007). Although isolation of pigs is sometimes necessary in management (e.g. during weighing or for medical treatments), management practices that keep an isolated pig within visual and/or olfactory range of its group, for example by using transparent or mesh partitions could reduce the chances of aggression when pigs are reintegrated into groups (Marchant & Marchant-Forde, 2005) (but see (Rushen, 1988)). After longer separations, facilitating the return to the social group in a step-wise process, for example by using a side compartment that provides opportunities for passive social buffering of separated pigs and assessment of cues by observers while initially preventing physical contact, may further reduce aggression and promote more prosocial interactions following reunions.

Our findings extend the growing evidence that pigs possess the cognitive and emotional capabilities necessary for empathic responses (Cordoni et al., 2023; Moscovice et al., 2023). Furthermore, they suggest that having opportunities for emotional contagion, while being able to control the timing and extent of social interactions with distressed partners, facilitates emotional regulation and the subsequent expression of consolation behaviour. However, our results also caution against attributing more general post-event increases in affiliation to consolation without carefully controlling for high-arousal confounds (Briefer, 2018; Pérez-Manrique & Gomila, 2018). This may be especially important to consider when studying consolation behaviour in herd species like pigs. Husbandry practices and experimental designs that allow graduated re-introductions of pigs who have experienced stressors may provide target pigs with opportunities for social buffering while group members have time to appraise the situation and potentially regulate their emotional responses. The combined effects may promote more prosocial behaviours in pigs. Our helping paradigm satisfies these conditions and therefore represents a promising framework for future research on a range of empathic behaviours in group-living animals.

