## Supplementary material for "Consolation behaviour in pigs: Prior exposure to group members in need of help drives targeted affiliation and facilitates social buffering"

**Supplementary Materials- Consolation behaviour in pigs**

**Supplementary video**

The supplementary video shows pig 7 being released into the home pen by a peer in the helping context, after which it receives affiliative contact consistent with consolation.

**Supplementary Methods**

**Table S1.** Ethogram used for behavioural coding. For interactions, we coded the initiator of the interaction in all cases, with the exception of nose-nose contact, due to difficulty in determining the initiator.

| **Behaviour Category** | **Description** |
| --- | --- |
| **Affiliative interactions** | |
| Nose-nose contact | Two pigs touch noses or snouts (non-directional) |
| Nose-body contact | A pig uses its nose to touch a body part of a target pig (excluding its nose and snout) |
| Pawing | A pig uses a paw to touch a body part of a target pig (excluding its nose and snout) |
| Head-over | A pig puts its head on the back of a target pig while both pigs are standing; often followed by rest in contact |
| Social rubbing | A pig rubs part of its body against the body of a target pig, while both pigs are standing. |
| **Non-Affiliative/Aggressive interactions** | |
| Lifting | A pig attempts to displace a target pig by lifting or levering it with snout or head |
| Mounting | A pig places its front legs across the back or rear of a target pig and attempts to rear up |
| Kicking | A pig projects one or both hind limbs towards a target pig and strikes it |
| Pushing | A pig presses its head, neck, shoulder or body against a target pig, causing it to move from the position that it was in |
| Chasing | A pig pursues a target pig, who walks or runs away |
| Head-knocking | A pig lurches or jerks its head hitting another pig |
| Fighting | Two pigs mutually push one another in a head-to-head orientation. The pattern can also involve body-to-body rotation and/or aggressive mounting, lifting, biting, attempt biting, kicking, chasing, head knocking and/or high-pitched vocalisations, lasting more than 10 s |
| **Anxiety behaviours** | |
| Body scratching/rubbing | A pig uses its legs or a substrate to rub part of its body |
| Vacuum-chewing | A pig chews with empty mouth |
| Head/body-shaking | A pig vigorously shakes its head and/or body |
| Yawning | A pig performs deep, long inhalation with open mouth |

**Supplementary Results**

**Table S2**. Results of a LMM testing for an effect of context (control, helping, direct-reunion) on total affiliation (including non-directional nose-nose contacts) between focal pigs and their group members. Significant predictors are indicated in bold.

| predictor | estimate | | s.e. | df | t or F value | *p value* |
| --- | --- | --- | --- | --- | --- | --- |
| (Intercept) | 0.898 | | 0.077 |  |  | (a) |
| **context_helping** | 0.484 | | 0.084 | 2, 210 | 72.614 | **< 0.001** |
| **context_direct reunion** | 1.009 | 0.084 | |  |  | (b) |
| sex_female | 0.016 | 0.07 | | 1, 210 | 0.225 | 0.822 |
| elo rank | -0.019 | 0.035 | | 1, 215 | -0.539 | 0.591 |

^a^ Not indicated due to limited interpretability of intercepts

^b^ Refer to F value and p-value for ‘context_helping’ for overall effect of predictor.

**Table S3.** Pairwise post hoc comparisons of estimated marginal means for the amount of total affiliation (including non-directional nose-nose contacts) that focal pigs are involved in across the different contexts. Significant differences are indicated in bold.

| contrast | estimate | s.e. | t ratio | adj. p val |
| --- | --- | --- | --- | --- |
| control - helping | -0.484 | 0.0838 | -5.779 | **< 0.001** |
| control - direct reunion | -1.009 | 0.0838 | -12.048 | **< 0.001** |
| helping - direct reunion | -0.525 | 0.0838 | -6.269 | **< 0.001** |

**Table S4**. Results of a LMM indicating that the interaction of context (control, helping, direct-reunion) and direction (give or get) has a tendency to influence the latency to affiliate between focal pigs and their group members.

| predictor | estimate | s.e. | df | F value | *p value* |
| --- | --- | --- | --- | --- | --- |
| (Intercept) | 8.375 | 0.56 |  |  | (a) |
| context_helping | -2.776 | 0.681 |  |  | (a) |
| context_direct-reunion | -4.684 | 0.667 |  |  | (a) |
| direction_give | -0.132 | 0.756 |  |  | (a) |
| sex_female | 0.241 | 0.397 | 1, 354 | 0.351 | 0.554 |
| elo rank | -0.076 | 0.202 | 1, 358 | 0.147 | 0.701 |
| context_direct-reunion:direction_give | -0.545 | 0.971 | 2, 353 | 2.407 | 0.092 |
| context_helping:direction_give | 1.443 | 1.016 |  |  | (b) |

^a^ Not indicated due to limited interpretability, either of intercepts or due to a tendency towards significant interaction terms

^b^ Refer to F value for ‘context_helping:direction_give’ for overall effect of interaction

**Table S5.** Pairwise post hoc comparisons of estimated marginal means for the latency for focal pigs to give and receive affiliation across the different contexts. Significant differences are indicated in bold.

| contrast | estimate | s.e. | t ratio | adj. p val |
| --- | --- | --- | --- | --- |
| direction = get |  |  |  |  |
| control – direct reunion | 4.68 | 0.675 | 6.944 | **<.0001** |
| control - helping | 2.78 | 0.689 | 4.029 | **0.0002** |
| direct reunion - helping | -1.91 | 0.631 | -3.024 | **0.0075** |
| direction = give |  |  |  |  |
| control - direct reunion | 5.23 | 0.713 | 7.333 | **<.0001** |
| control - helping | 1.33 | 0.762 | 1.749 | 0.1886 |
| direct reunion - helping | -3.9 | 0.67 | -5.813 | **<.0001** |

**Table S6.** Pairwise post hoc comparisons of estimated marginal means for the amount of directional affiliation that focal pigs give and get across the different contexts. Significant differences are indicated in bold.

| contrast | estimate | s.e. | z ratio | adj. p val |
| --- | --- | --- | --- | --- |
| direction = get |  |  |  |  |
| control - helping | 0.439 | 0.053 | -6.771 | **< 0.001** |
| control - direct reunion | 0.286 | 0.034 | -10.584 | **< 0.001** |
| helping - direct reunion | 1.534 | 0.160 | -4.097 | **< 0.001** |
| direction = give |  |  |  |  |
| control - helping | 0.884 | 0.042 | -0.924 | 0.627 |
| control - direct reunion | 0.349 | 0.042 | -8.634 | **< 0.001** |
| helping - direct reunion | 2.534 | 0.300 | -7.854 | **< 0.001** |

**Table S7.** Pairwise post hoc comparisons of estimated marginal means for the total amount of non-affiliative interactions (given or received) that focal pigs are involved in across the different contexts. Significant differences are indicated in bold.

| contrast | estimate | s.e. | t ratio | adj. p val |
| --- | --- | --- | --- | --- |
| control / direct reunion | 0.493 | 0.090 | -3.872 | **< 0.001** |
| control / helping | 0.747 | 0.140 | -1.554 | 0.2658 |
| direct reunion / helping | 1.515 | 0.263 | 2.395 | **0.0438** |

**Table S8.** Pairwise post hoc comparisons of estimated marginal means (odds ratios) for likelihood of showing anxiety across the three different contexts. Significant differences are indicated in bold.

| contrast | odds ratio | s.e. | z ratio | p value |
| --- | --- | --- | --- | --- |
| control / direct reunion | 0.55 | 0.263 | -1.252 | 0.4228 |
| control / helping | 1.72 | 0.758 | 1.231 | 0.4348 |
| direct reunion / helping | 3.13 | 1.41 | 2.531 | **0.0305** |

**Table S9.** Results of a LMM indicating a significant effect of expression of anxiety behaviours on changes in salivary cortisol concentrations in focal pigs (from pre- to post-test) during helping and direct-reunions. The impact of affiliative and non-affiliative behaviours received on overall cortisol concentrations (both pre- and post-test) is also shown. Significant effects are indicated in bold..

| predictor | est. | s.e. | df | t or F value | *p- value* |
| --- | --- | --- | --- | --- | --- |
| (Intercept) | 2.083 | 0.079 |  |  | (a) |
| affiliation received | -0.014 | 0.031 | 1, 255 | -0.464 | 0.636 |
| non-affiliative received | 0.058 | 0.029 | 1, 265 | 1.991 | 0.051 |
| context_helping | -0.142 | 0.103 | 1, 267 | -1.386 | 0.170 |
| condition_pre-test sample | -0.533 | 0.063 |  |  | (a) |
| anxiety_yes | 0.032 | 0.082 |  |  | (a) |
| sex_female | -0.035 | 0.071 | 1, 65 | -0.492 | 0.639 |
| elo rank | 0.002 | 0.036 | 1, 68 | 0.069 | 0.944 |
| test order | -0.041 | 0.035 | 1, 75 | -1.183 | 0.259 |
| sample collection time | -0.004 | 0.035 | 1, 240 | -0.11 | 0.905 |
| duration stressor to sample | -0.012 | 0.051 | 1, 261 | -0.236 | 0.797 |
| **condition_pre:anxiety_yes** | 0.225 | 0.098 | 1, 205 | 5.106 | **0.025** |

^a^ Not indicated due to limited interpretability, either of intercepts or due to significant interaction terms

**Table S10.** Pairwise post hoc comparisons of estimated marginal means for the interaction effect of sample (pre vs. post experiment) and indicators of anxiety (yes or no) on cortisol concentrations. Significant differences are indicated in bold.

| contrast | estimate | s.e. | t ratio | adj. p val |
| --- | --- | --- | --- | --- |
| Pre sample |  |  |  |  |
| Anxiety 0 / 1 | 0.773 | 0.0638 | -3.122 | **< 0.001** |
| Post sample |  |  |  |  |
| Anxiety 0 / 1 | 0.968 | 0.0814 | -0.382 | 0.7024 |

**Supplementary Figure S1.** Changes in salivary cortisol concentrations in the helping and direct-reunion contexts.

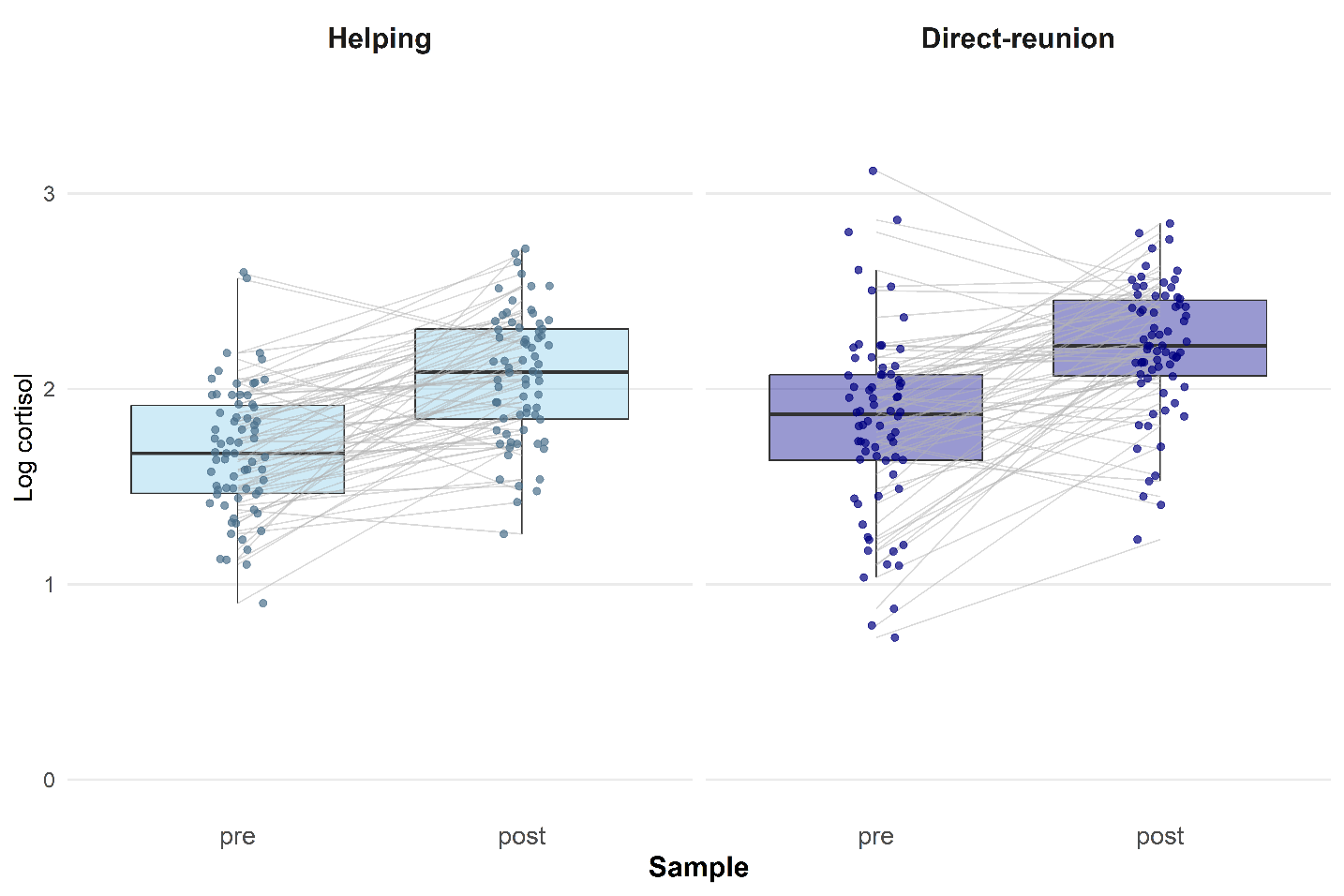
